# Resilience trinity: safeguarding ecosystem services across three different time horizons and decision contexts

**DOI:** 10.1101/549873

**Authors:** H Weise, H Auge, C Baessler, I Bärlund, E.M. Bennett, U Berger, F Bohn, A Bonn, D Borchardt, F Brand, A Chatzinotas, R Corstanje, F De Laender, P Dietrich, S Dunker, W Durka, I Fazey, J Groeneveld, CSE Guilbaud, H Harms, S Harpole, J Harris, K Jax, F Jeltsch, K Johst, J Joshi, S Klotz, I Kühn, C Kuhlicke, B Müller, V Radchuk, H Reuter, K Rinke, M Schmitt-Jansen, R Seppelt, A Singer, RJ Standish, HH Thulke, B Tietjen, M Weitere, C Wirth, C Wolf, V Grimm

## Abstract

Ensuring ecosystem resilience is an intuitive approach to safeguard future provisioning of ecosystem services (ES). However, resilience is an ambiguous concept and difficult to operationalize. Focusing on resilience mechanisms, such as diversity, network architectures or adaptive capacity, has recently been suggested as means to operationalize resilience. Still, the focus on mechanisms is not specific enough because the usefulness of a mechanism is context-dependent. We suggest a conceptual framework, *resilience trinity,* to facilitate management of resilience mechanisms in three distinctive decision contexts and time-horizons. i) reactive, when there is an imminent threat to ES resilience and a high pressure to act, ii) adjustive, when the threat is known in general but there is still time to adapt management, and iii) provident when time horizons are very long and the nature of the threats is uncertain, leading to a low willingness to act. This emphasizes that resilience has different interpretations and implications at different time horizons which however need to be reconciled. The inclusion of time into resilience thinking ensures that longer-term management actions are not missed while urgent threats to ES are given priority.

## 1. Introduction

Resilience is a characteristic feature of ecosystems, otherwise they would not exist. It describes their ability to resist, or recover from, disturbances and to maintain functioning (Oliver *et al.* 2015). Losing resilience of ecosystems might put their continued provision of ecosystem functions and services (ES) at risk. Thus, understanding resilience of ecosystems, and its limits, is of fundamental interest because humans depend on ecosystem services (Millennium Ecosystem Assessment 2005; Díaz *et al.* 2015). Management for sustainable ES provisioning must safeguard, strengthen, or restore ecosystems’ resilience. However, the utilization of these insights in practice is still limited. While it makes intuitive sense to manage for resilience it is unclear which actions should follow from this goal (Standish *et al.* 2014).

One reason that hampers managing for resilience is the broad use of the concept itself. Current interpretations range from resilience as a way of thinking in sustainability science (Folke *et al.* 2010; Biggs *et al.* 2015) to resilience as a multidimensional metric comprising recovery, resistance, and persistence in ecology and biodiversity research (Oliver *et al.* 2015; Donohue *et al.* 2016; Ingrisch and Bahn 2018), through to adopting resilience as a management paradigm in response to national policy (e.g. Isaac et al, 2018). However, if resilience is to be operationalized, this broad range of interpretations creates at least confusion, many cases of false labelling (Donohue *et al.* 2016), and at worst loopholes for mismanagement (Schoon *et al.* 2015; Newton 2016). Likewise, quantification of resilience (Angeler and Allen 2016; Allen *et al.* 2016) will remain a major issue unless the multiple meanings and implications of resilience have not been disentangled. How to move on despite these difficulties?

As a way forward it has been suggested to focus on managing specific mechanisms that underlie the resilience of ecosystem functioning (Biggs *et al.* 2012; Oliver *et al.* 2015; Berthet, et al, 2018) instead of focusing on managing for resilience *per se*. Resilience mechanisms are the mechanisms underlying recovery, resistance, and persistence. Focusing on mechanisms helps us to be more specific about which outcome exactly we want to be resilient and about the concrete steps to achieve this. We argue, however, that a sole focus on resilience mechanisms is still not specific enough.

Consider an example from forestry: bark beetle infestations can kill off entire stands of spruce and cause great damage (Wermelinger 2004). Thus, the ES of wood provision is strongly reduced in a short-term perspective. A countermeasure could be spruce reforestation, where thinning would aim at increasing the fitness of individual trees and thus focus on resilience mechanisms at the level of the individuals (Seidl *et al.* 2016). However, if we consider a longer time horizon (e.g. centuries), the effects of insect outbreaks in temperate European forests may be exacerbated by other disturbances, such as water limitation, forest fires or outbreaks of tree-killing pathogens (Lindner *et al.* 2010; Seidl *et al.* 2016). On that longer time frame, just spruce reforestation and thinning would not be the most useful way to strengthen the resilience of wood production. Instead, fostering resilience mechanisms at the community level, for example by increasing stand heterogeneity, would be a better choice because interspecific differences in reactions to disturbances can be utilized to ensure long term wood supply. On the other hand, it will take longer to establish a more diverse forest than a monoculture, which implies short-term economic losses that might not be tolerable.

How should we deal with these kinds of trade-offs? Approaches are needed that account for our limited understanding of system responses while advocating the use of natural mechanisms. Here, we suggest a conceptual framework that comprises resilience mechanisms and time horizons to facilitate better decisions to safeguard future ecosystem service provisioning. In this article, we (1) provide a brief review and a new categorization of resilience mechanisms, (2) suggest three time horizons for the management of ecosystem services and show that they imply different decision contexts, (3) give an example on the linkage of time horizons and resilience mechanisms, and (4) discuss how our framework could be embedded in environmental decision making. Our key message is that resilience is not one thing but has different interpretations in different decision contexts (“*resilience trinity*”).

## 2. Resilience mechanisms

Resilience mechanisms have been explored for decades in systems as diverse as coral reefs, rangelands, rainforests, or contaminated aquifers. Table 1 provides an overview of mechanisms that have been identified in review articles focusing on “resilience”, “mechanism”, and “ecosystem service” (see Supporting Information for the specific definition of each mechanism). We grouped the mechanisms into three categories. (1) *Portfolio* mechanisms spread the risk of being affected by a disturbance. They are often based on diversity, redundancy, or heterogeneity. Mechanisms in the category *Function* (2) are related to important roles that elements of a system play for functioning; they can only be observed dynamically as they enfold in the course of time. A well-known mechanism in this category is the presence of keystone species. Some overlap with the portfolio category exists. However, here the focus is on functional aspects that are not primarily based on diversity, redundancy, or heterogeneity. (3) *Adaption* mechanisms share aspects of the *Portfolio* and *Function* category. They require diversity to function and are observed over the course of time. However, resilience mechanisms in this category are different because they feature adaptation via various mechanisms, including natural selection. (4) The fourth category, *Structure,* refers to structural features that affect recovery and resistance and that can be observed statically, in a snapshot of a system. Prominent examples are modularity and connectivity. The main purpose of Table 1 is to demonstrate the diversity of resilience mechanisms and that any attempt to categorize them is necessarily subjective and to some degree arbitrary, as can be inferred from comparing our categories to those cited in the legend of Table 1 and the supplement. The reason is that most mechanisms do not work in isolation but together with other mechanisms, but whether and how they do so depends on the specific system and context under consideration.

**Table 1.** Resilience mechanisms. So far, no systematic or comprehensive overview of resilience mechanisms exists. Here we compiled ecological mechanisms from various reviews and grouped them into categories (see Table S1 for definitions of each mechanism). Those categories are not exclusive. Other possible categories are for example diversity, connectivity and adaptive capacity (Bernhardt and Leslie 2013); species, community, landscape (Oliver et al. 2015); complexity, adaptivity (Desjardins et al. 2015). For a comprehensive overview of mechanisms in social-ecological systems see (Biggs et al. 2012, 2015), and for attributes that confer resilience to climate change in the context of restoration see Timpane-Padgham et al. (2017).

| GROUP | MECHANISM | GENERAL IDEA | EXAMPLE DEFINITIONS | REFERENCES |
| --- | --- | --- | --- | --- |
| *Portfolio:*<br>*Spreading the*<br>*effects of*<br>*disturbances* | **Redundancy**<br>(functional,<br>species) | Losing certain species may not<br>matter because their function can<br>be provided by functionally<br>equivalent species. | “when multiple species perform similar<br>functions (...) the resistance of an ecosystem<br>function will be higher if those species also<br>have differing responses to environmental<br>perturbations”(Oliver et al. 2015) | (Biggs *et al.* 2012; Bernhardt and Leslie 2013;<br>Griffiths and Philippot 2013; Desjardins *et al.*<br>2015; Oliver *et al.* 2015; Sasaki *et al.* 2015) |
|  | **Diversity**<br>(genetic, habitat,<br>species, trait,<br>response) | Individuals or populations are<br>sensitive to disturbances to<br>different extents. | “Species in the same functional group often<br>show different responses to disturbances<br>(Laliberte et al. 2010), and hence the value of<br>redundancy” | (Palumbi *et al.* 2009; Chapin III *et al.* 2010;<br>Traill *et al.* 2010; Biggs *et al.* 2012; Bernhardt<br>and Leslie 2013; Griffiths and Philippot 2013;<br>Thompson *et al.* 2014; Desjardins *et al.* 2015;<br>Oliver *et al.* 2015; Sasaki *et al.* 2015) |
|  | **Heterogeneity**<br>(stand,<br>landscape) | Functions lost in certain places<br>can be compensated or restored<br>from other, less affected places. | “Resilience is an emergent ecosystem property<br>conferred through biodiversity, (...),<br>ecosystem diversity (heterogeneity and beta<br>diversity) across a forest landscape“<br>(Thompson et al. 2014) | (Thompson *et al.* 2014; Desjardins *et al.* 2015) |
|  | **Area of habitat**<br>**cover at the**<br>**landscape scale** |  | “Larger areas of natural or semi-natural habitat<br>tend to provide a greater range and amount of<br>resources, which promote higher species<br>richness and larger population sizes [...]. This<br>[...] is likely to mean greater genetic diversity<br>and functional redundancy [...]” (Oliver et al.<br>2015) | (Oliver *et al.* 2015) |
| **Function:**<br>*Functional*<br>*features that*<br>*affect recovery*<br>*and resistance*<br>*(identifiable*<br>*only in system* | **Negative**<br>**feedbacks** | Negative feedbacks, for example<br>density dependence, cause<br>recovery to equilibria. | “Negative feedback mechanisms contribute to<br>maintain the ecosystem state” (Conversi et al.<br>2014) | (Chapin III *et al.* 2010; Gedan *et al.* 2011; Biggs<br>*et al.* 2012; Conversi *et al.* 2015; Spears *et al.*<br>2015) |
|  | **Keystone**<br>**species** | Keystone species (of functional<br>types) may be the main factor | “Loss of the keystone species can lead to<br>cascading effects” (Sasaki et al. 2015) | (Traill *et al.* 2010; Thompson *et al.* 2014; Sasaki<br>*et al.* 2015) |
| *dynamics)* |  | maintaining a certain function or structure. | “Loss of keystone predators can have large effects for a system through cascading effects of expansion of herbivore populations” (Thompson et al. 2014) |  |
|  | **Dominant species** | A dominant species that is resilient will entail its resilience to the entire system. | “If the dominant species is resilient to disturbances it will maintain ES functioning despite disturbances” (Sasaki et al. 2015) | (Sasaki *et al.* 2015) |
|  | **Strength of species interaction** | Weak links in an interaction network of species can dampen internal and external variations in components of the network. | “Weakly interacting species stabilize community dynamics by dampening strong, potentially destabilizing consumer-resource interactions and facilitative interactions.” (Bernhardt & Leslie 2013) | (Bernhardt and Leslie 2013) |
|  | **Separation of time scales (“slow variables”)** | Through self-organization, slow-changing variables can emerge that constrain and control fast-changing variables and thereby reduce variation (“panarchy”) | “slow variables are usually related to regulating ecosystem services, and that the strength of regulating services can attenuate the impact of shocks on ecosystems.” (Bennett et al. 2009) | (Bennett *et al.* 2009; Biggs *et al.* 2012) |
| **Adaptation:**<br>*Adaptation in order to better* | **Adaptive phenotypic plasticity** | Individuals change in response to disturbance and thereby reduce the effects of subsequent | “Capacity of individuals to respond to environmental changes through flexible behavioural or physiological strategies [...]” | (Bernhardt and Leslie 2013; Oliver *et al.* 2015) |
| *cope with disturbances; changes in disturbance regimes* |  | disturbances. | (Oliver et al. 2015)<br>“(...) phenotypic plasticity may be the most important component of adaptive potential (...). ” (Bernhardt & Leslie 2013) |  |
|  | **Adaptive capacity** | Populations, communities, and ecosystems have the ability to change, through change of individuals, shifts of distributions, and rapid evolution, and thereby reduce responses to subsequent disturbances. | “ability of populations, communities and ecosystems to adapt [...] through a combination of phenotypic plasticity, physiol. responses, distributional shifts, rapid evolution of traits” (Bernhard and Leslie 2013) | (Bernhardt and Leslie 2013) |
|  | **Learning** | Individuals learn and thereby recover faster or respond less to disturbances. | “The process of modifying existing or acquiring new knowledge, behaviours, skills, values, or preferences at individual, group, or societal levels”(Biggs et al. 2012) | (Biggs *et al.* 2012, 2015) |
| *Structure: Structural features that affect recovery* | **Connectivity** | Higher connectivity between habitats allows for faster recovery by moving individuals or resources. | “connections that promote stability and recovery at multiple scales of biological organization” (Bernhardt and Leslie 2013) | (Biggs *et al.* 2012; Bernhardt and Leslie 2013) |
| *and resistance*<br>*(identifiable in snapshots of a system)* | **Modularity** | Organization in more or less disconnected compartment allows for asynchronous responses and thereby recovery. | “It refers to compartmentalization of populations in space and time. (...). For example, where populations are too closely connected, severe disturbances to one population may affect all populations.”<br>(Bernhardt & Leslie 2013) | (Bernhardt and Leslie 2013) |
|  | **Network architecture** | Depending on the type of interaction between nodes, the connectedness and other features of the interaction network determine the response to disturbances. | “A highly connected and nested architecture promotes community stability in mutualistic networks, whereas stability is increased in compartmented and weakly connected architectures in trophic networks. (...)”<br>(Bernhardt & Leslie 2013) | (Bernhardt and Leslie 2013; Griffiths and Philippot 2013; Oliver *et al.* 2015) |
|  | **Spatial self-organization** | Positive feedback can lead to self-organized spatial patterns that are self-similar over time and lead to recovery from disturbances. |  |  |

The mechanisms listed in Table 1 represent empirical knowledge, theory, or expert knowledge. However, it seems impossible to translate the knowledge they represent directly into actions. For example, intuitively it seems evident that biodiversity increases resilience, but decades of biodiversity research show how difficult it is to prove and understand this relationship in systems that are more complex than simplified models or controlled experiments (Cardinale *et al.* 2012; De Laender *et al.* 2016). Thus, the mechanisms listed in Table 1 are only possible mechanisms. Whether they are relevant, whether some of them dominate or compromise others depends on the specific situation (Biggs et al. 2012; Desjardins et al. 2015).

## 3. Rationale for a “resilience trinity”

Addressing the resilience of ecosystem services and, hence, functioning, forces us to ask specific questions: what specific services and potential disturbances are we talking about? That is, we have to answer the question “resilience of what to what” (Carpenter *et al.* 2001). The disadvantage is that the concept of ES itself is subject of critical debate. Issues include trade-offs between ES (Seppelt *et al.* 2011), the delineation of “ES providing units”, and whether or not biodiversity is a service, a good, or a mechanism (Bennett *et al.* 2009; Mace *et al.* 2012; Jax and Heink 2015).

The focus on ES nevertheless helps us to address specific resilience mechanisms. In contrast to managing the resilience of a hard-to-define ecosystem, the relevant level of biological organization often is obvious, if the goal is to manage for resilience of a specific ES. However, clarity about the relevant ES and ecosystem functions is still not sufficient. Consider for example the storage of organic carbon in soils (soil organic carbon, SOC). Soils store at least three times the amount of carbon found in either the atmosphere or in living plants (Parry *et al.* 2007). This is one key function supporting climate regulation. Soil biota mediate SOC persistence and turnover (Schmidt *et al.* 2011; Schimel and Schaeffer 2012). Land use is one of the main stresses on SOC levels, with persistently low and decreasing levels in more intense land uses. Stress leads to the destruction of soil structure and to a decrease in soil biodiversity on which structural reformation relies (Crawford *et al.* 2011; Ponge *et al.* 2013). Short term (∼1yr) responses to conserve SOC aim at improving soil and land management practices, such as less intense tillage, retaining stubble in the field, or introducing cover crops when fields are temporarily not in production. Longer term (10-100 yr) measures would comprise similar soil management interventions but also consider taking fields out of production permanently. Additional options are the introduction of intercropping systems or even landscape engineering, for example producing terraces. This means to consider long-term land use and its management practices from a planning and policy perspective, for instance designating land-uses (e.g., forestry) on lands prone to SOC loss.

Thus, different time horizons require different measures. Threats to ES can be acute and obvious in some contexts. In these situations, the loss of the desired functions is imminent or has already happened. Time for *reaction* is limited and the actions are planned for comparatively short time horizons. We call this decision context *reactive*. It is further characterized by a high acceptance for actions by the stakeholders involved. Examples for a *reactive decision context* include local pest outbreaks, an emerging wildlife disease that threatens livestock, or catastrophic floods in river flood plains. The “command and control” mindset of engineering usually is dominant in this context.

In contrast to *reactive*, in *adjustive* decision contexts ES are threatened, but not yet to a level that is critical to their provisioning. Concerns about future losses exist, but the perceived urgency of actions to increase resilience is lower than in a *reactive* context. Therefore, there are initiatives and incentives to *adjust* current management practices. Safeguarding ES resilience in an *adjustive* decision context can be slow though or even fail because the lower perceived urgency for actions. An example for an *adjustive* decision context is the safeguarding of pollination services, for example by trying to revert the decline of wild bee and other pollinator populations. Mostly, this is the context in which typically ecologists discuss resilience.

Third, *provident* decision contexts are distinguished from the two previous contexts by even longer time horizons. Here, the task is to conserve, restore or improve resilience mechanisms without a specific threat being the trigger for actions. The basic motivation is that currently environmental and societal drivers are changing at unprecedented rates and might change even faster in the future. These changes might “erode” the resilience of ES in all kinds of unforeseeable ways. The basic motivation to safeguard ES resilience at these longer time horizons is the insight that if we carry on management as before, we will lose ES. However, we do not know how, when and because of which specific threats they will be lost. Lacking the imminent threat, acceptance of actions is low for provident decision contexts, particularly because returns from current investments are uncertain. An example is the creation of large reserve networks, which can safeguard nature even against unknown future threats, as they are likely to harbour the structure and functions required for resilience mechanisms with high capacity. However, this benefit cannot easily be accounted for and thus not be balanced against the loss of, for example, arable land. Typically, this provident decision context is addressed under the umbrella sustainability or transformation.

Our idea of provident resilience is similar to the idea of “general resilience” (Folke et al. 2010), which is “concerned more about resilience to all kinds of shocks, including completely novel ones” (Folke et al. 2010), while their “specified resilience” refers “to problems relating to particular aspects of a system that might arise from a particular set of sources or shocks.” (Folke et al. 2010). Our “resilience trinity” framework more explicitly refers to different time horizons and decision contexts, but overlaps with the specified/general distinction by emphasizing the long-term risks of focussing solely on reactive, or specified, resilience.

The *resilience trinity* framework refers to the threefold potential of reactive, adjustive, and provident decision contexts for managing the resilience of ES. In the following, we will give an example by applying our conceptual framework to the ES of water purification (Fig. 1).

**Figure 1.**
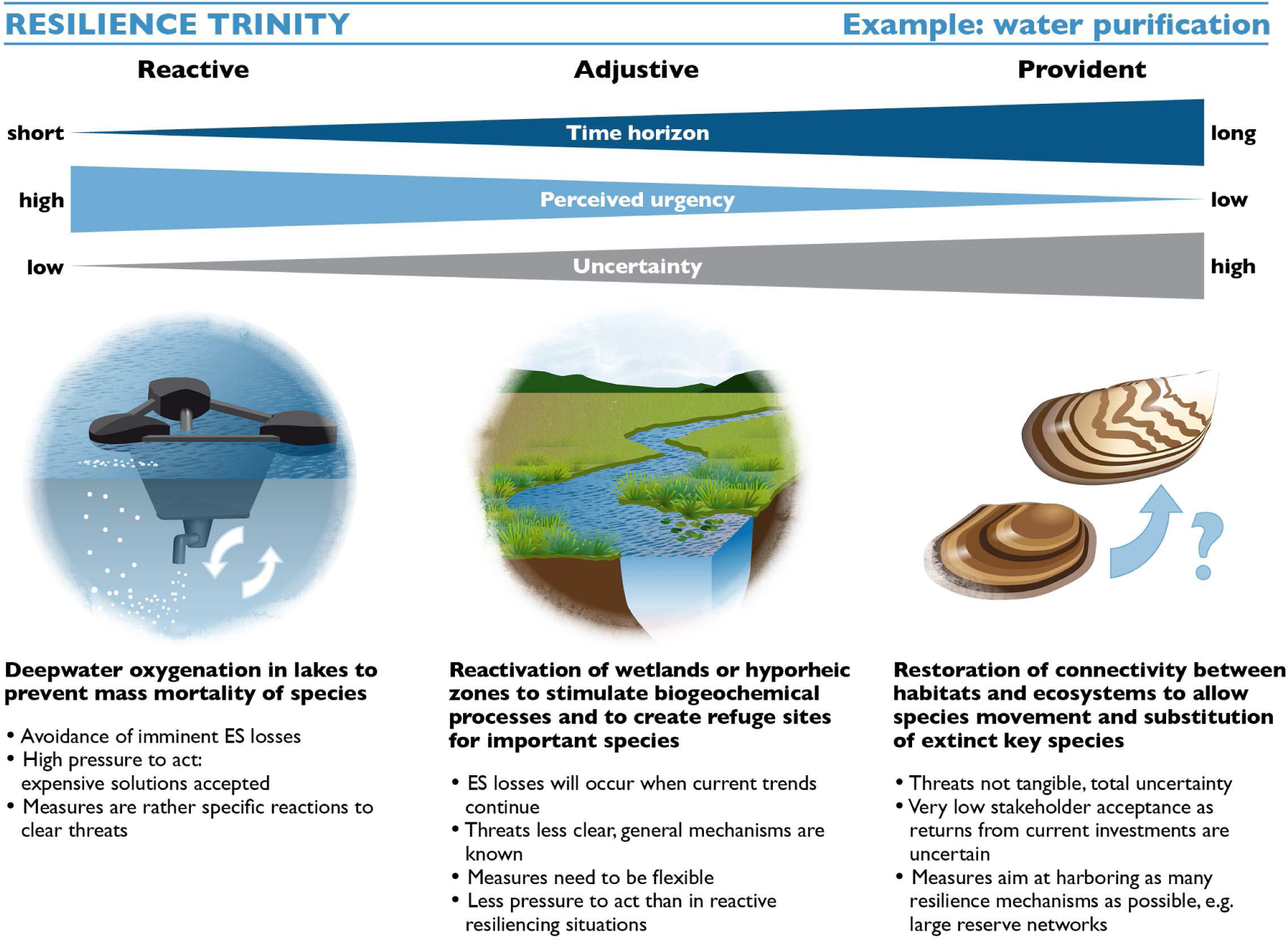
Measures to safeguard the ecosystem service (ES) of water purification across different time horizons. The time horizon of interest determines the decision context (upper arrow). If an ES needs to be safeguarded now (very short time horizon) the pressure to act is high and uncertainty is comparatively low. In contrast, uncertainty is very high and the pressure to act very low for long time horizons. The resulting decision contexts will warrant different measures; thus we propose to distinguish *reactive, adjustive* and *provident* contexts. It is important to be aware of the different decision contexts because they will lead to different decisions and trade-offs. Our resilience trinity framework tries to create and establish this awareness.

## 4. Example: Water purification

Water is a fundamental resource. Societies, economies and the natural environment rely on permanent water provision in sufficient quantity and quality. A multitude of threats endanger water purification services. Some threats are acute (e.g., pulses of toxicants or the occurrence of extensive anoxia in lakes) and require immediate action. At intermediate time scales, solutions must be found to control pollution pathways, avoid structural degradation of river courses or excessive eutrophication. On longer time scales, threats are likely related to human perturbations of the global biogeochemical cycles (nitrogen, phosphorus, carbon), predominantly by farming, or structural degradations. However, direct and indirect consequences from these threats cannot be forecasted yet, which makes it more challenging to develop and justify countermeasures.

Actions related to *reactive* decision contexts aim to prevent the imminent danger of losing ecosystem functions. They are often based on technology (Fig. 1), such as the oxygenation of deep water in lakes to prevent mass mortality of species and internal loading with pollutants (Beutel and Horne 1999). These measures profit from extensive knowledge of the ecosystem’s functioning and a clear definition of the problem and its solution. High expenses of these solutions are justifiable by the high societal pressure to act, e.g. for maintaining drinking water supply from a reservoir or a lake (e.g. suppression of manganese release, Bryant et al. (2011)). In addition, they are a reaction to a specific and rather clearly described threat.

An example for an action in an *adjustive* decision context is the nutrient reduction by flocculation, which is used to remove nutrients from lakes or reservoirs (Mehner *et al.* 2008) and add substances such as aluminium to remove phosphate from the water column. Lowering phosphate concentrations reduces eutrophication in general and shifts algal communities from a dominance of potentially toxic cyanobacteria towards a community consisting of eukaryotic algae and therefore comes along with a major improvement of water quality. Such measures require more careful planning than *reactive* decisions, for example adapted dosing and application of flocculants, as well as detailed pre-studies. Another example of a decision in an *adjustive context* is the activation of major reactive zones, for example wetlands or hyporheic zones in riverbeds (Rode *et al.* 2015) for water quality regulation.

Decisions in *provident* contexts often follow a systems approach. In the water sector they often include the landscape context. Implementation of buffer strips along rivers, for example, can reduce nutrient exports from land into the water cycle and therefore weaken environmental pressures from agriculture (Mayer *et al.* 2007). Another example are key species, which are often important for self-purification within aquatic environment such as bivalves filtrating the water or other organisms with similar functions (McCay *et al.* 2003; Kathol *et al.* 2011). The protection of key species is one example for safeguarding ES resilience at long time horizons. This includes the restoration of habitats and refuges for key species. But still, current key species may not prevail under future conditions. Thus, actions that allow other species with the same functions but a better fitness to thrive under new conditions could be an important measure in *provident* decision contexts. Managing connectivity between ecosystems and habitats to allow for spread of better adapted species while monitoring the effects of species’ movements on the functionality of food webs could thus be a core element to safeguard the resilience of water purification at long time horizons. In that sense, invading species may even substitute the loss of natural key species. In the future, invading mussels and clams could substitute native unionids in rivers of the Northern Hemisphere, a pattern which is already present in several systems (Strayer and Smith 1996; Caraco *et al.* 2006). This may, however, be at odds with various other goals such as biodiversity conservation, maintaining existing ecosystem functions, functionality of human infrastructures or human recreation (Minchin *et al.* 2002). Thus, the action must be discussed with many different stakeholders.

## 5. How to integrate our framework in environmental decision making?

To elaborate how the resilience trinity framework could possibly support decision-making, we illustrate its application in combination with scenario planning. Scenario planning, such as formative scenario analysis, is an established tool in environmental decision making to conceptualize the future in a structured way (Peterson *et al.* 2003; Polasky *et al.* 2011; Brand *et al.* 2013). We suggest an iterative process facilitating the identification of relevant decision contexts and ecological mechanisms (Fig. 2). This requires the setting of goals for ES provisioning (upper panel of Fig. 2) and the identification of trade-offs, for example between different bundles of ecosystem services and between the interests of different groups (Schoon *et al.* 2015). Deliberating goals for ES is a political process which requires debates on intra and inter-generation fairness, on economic development pathways and on substituting ES through technologies (Jax *et al.* 2013) and thus negotiations among diverse groups of stakeholders. When goals are defined, our framework can help to ask questions that guide towards the right actions (Fig. 2, Step A - C). Answering these questions will lead to more clarity about the ecological situation, the relevant resilience mechanisms, and about the threats to these mechanisms (Fig. 2, Step A, B); ecological situations are defined by the state variables, reference states, disturbances, and scales to be considered (Grimm and Wissel 1997). The scenarios should then be confronted with a “reality check” (Fig. 2, Step C): when scenarios exist, the costs of safeguarding ES resilience (e.g., trade-offs between current and future ES supply) become tangible and strategic decisions on the best scenario path must be made.

**Figure 2.**
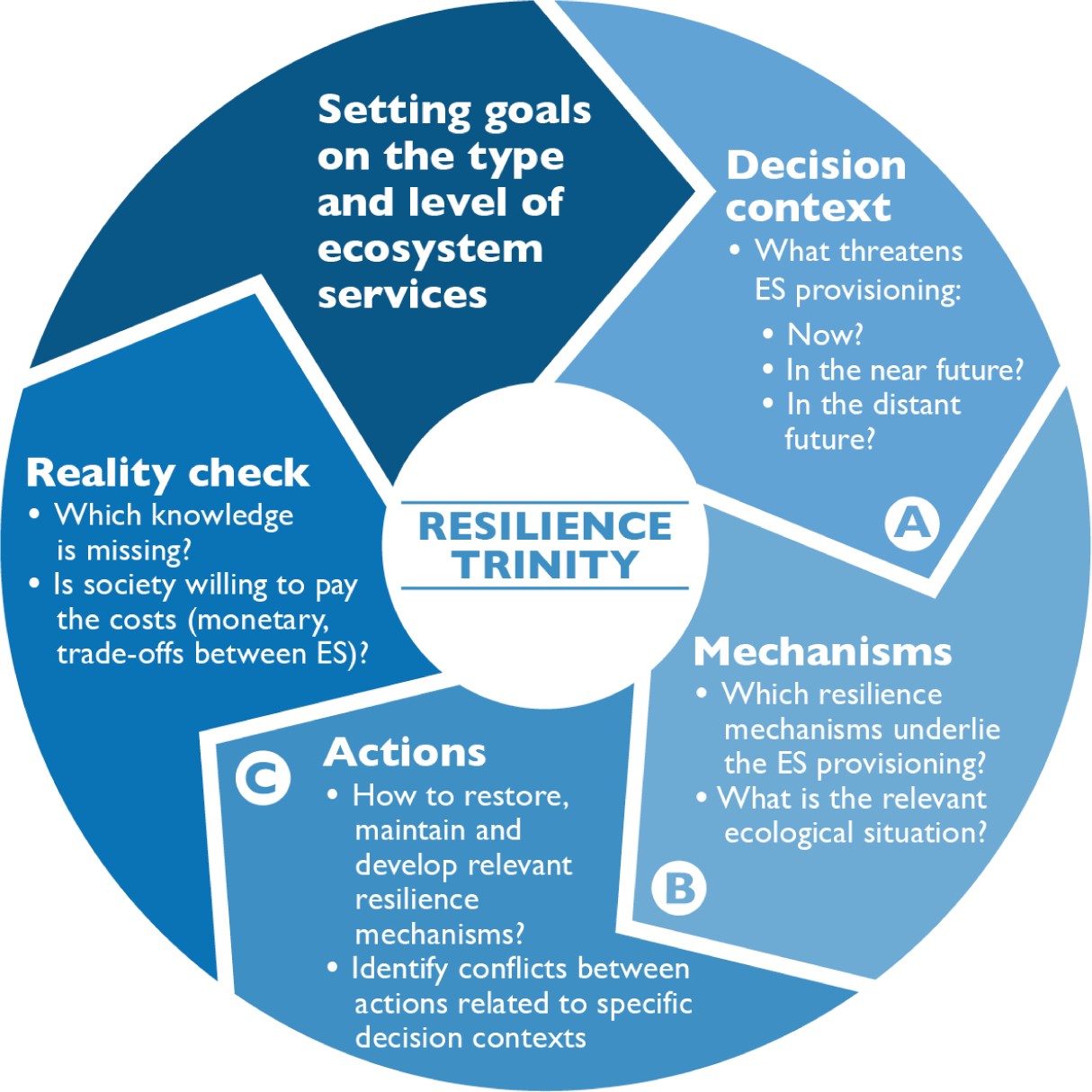
The use of the resilience trinity framework is to guide the identification of suitable actions to ensure sustained ES provisioning to society. The desired type and level of future ecosystem service provisioning needs to be decided through consultation. We suggest asking which resilience mechanisms are underlying the ecosystem service provisioning and to describe the ecological situation in which they apply. The next steps are the identification of threats to the resilience mechanisms and the clarification of the time horizon and thus the decision context. Step C is to identify actions to safeguard the resilience mechanisms. In addition, an analysis to assess potential conflicts between the actions will identify perverse outcomes. Steps A, B and C are not necessarily consecutive. Some iterative stepping back and forth may be necessary during the development of the scenarios. These scenarios will need to be assessed for feasibility given the willingness of society to pay for ES provisioning.

## 6. Discussion and Conclusions

Resilience is an important concept, both because it refers to essential features of ecosystems (resistance, recovery and persistence) and because it is a boundary concept that is popular among scientists, actors and stakeholders alike. However, the ambiguity of this concept has so far hampered its use it for planning, management, and environmental decision making (Standish *et al.* 2014).

Our framework, dubbed *resilience trinity*, tries to keep the attractiveness of the concept while demanding more specification. Our main purpose is to create awareness for different time horizons (short, intermediate and long-term). These imply different decision contexts and management attitudes (*reactive, adjustive and provident*). None of the three contexts is more important than the others – they all need to be considered and finally to be reconciled. Otherwise, exclusively focusing on reactive management could compromise long-term resilience of certain services. Solely provident actions could lead to short-term losses of services, or unacceptable risks. There is no simple, generic solution to reconcile the trade-offs of the different time horizons and decision contexts. Rather, our framework is designed to add structure to decision making and policy development (Fig. 2). Further examples and, preferably, case studies will be needed to learn about, and possibly improve, its usefulness.

A main criterion for the design of the *resilience trinity* framework was simplicity: focusing on three time horizons and clarifying the decision context can facilitate operationalizing resilience and ecosystem services, both complex and multidimensional concepts. Our vision is to ultimately foster a proactive approach to think about resilience: away from an absolute and desirable state towards a process with concrete actions. To foster this, in analogy to *stability-stabilizing* we might think about *resiliencing* ecosystem services. This has the advantage that we need to be specific: What is it that we want to *resilience*? And how exactly do we want to do it? The hope is that this approach will lead to a more action-oriented attitude towards resilience and thus put the concept to use for the safeguarding of future ecosystem service provisioning.

## Acknowledgments

We would like to thank Sebastian Fiedler, Hans-Jörg Vogel, Steve Railsback, and Graeme Cumming for helpful comments on an earlier version of this manuscript. HW acknowledges funding support through the German Research Foundation DFG project TI 824/2-1 “Ecosystem resilience towards climate change – the role of interacting buffer mechanisms in Mediterranean-type ecosystems and through the project Emerging Ecosystems”. We also thank UFZ’s Integrated Project “Emerging Ecosystems” within the research program “Terrestrial Environment” of the Helmholtz Association for funding workshops to develop ideas that are presented in this manuscript. We thank Lisa Vogel and Daphne Braun for professional revision of the illustrations.

## References

Angeler DG, Allen CR. 2016. Quantifying resilience. Journal of Applied Ecology 53: 617–624.

Allen CR, Angeler DG, Cumming GS, Folke C, Twidwell D, Uden DR. 2016. Quantifying spatial resilience. Journal of Applied Ecology 53: 625–635.

Bennett EM, Peterson GD, and Gordon LJ. 2009. Understanding relationships among multiple ecosystem services. Ecology letters 12: 1394–1404.

Bernhardt JR and Leslie HM. 2013. Resilience to climate change in coastal marine ecosystems. Annual Review of Marine Systems 5: 371–92.

Berthet, E.T., Bretagnolle, V., Lavorel, S., Sabatier, R., Tichit, M., and Segrestin, B. (2018) Applying ecological knowledge to the innovative design of sustainable agroecosystems. Journal of Applied Ecology doi 10.111/1365-2664.13173.

Beutel MW and Horne AJ. 1999. A review of the effects of hypolimnetic oxygenation on lake and reservoir water quality. Lake and Reservoir Management 15: 285–297.

Biggs R, Schlüter M, Biggs D, et al. 2012. Toward principles for enhancing the resilience of ecosystem services. Annual review of environment and resources 37: 421–448.

Biggs R, Schlüter M, and Schoon ML. 2015. Principles for building resilience: sustaining ecosystem services in social-ecological systems. Cambridge University Press.

Brand FS, Seidl R, Le QB, et al. 2013. Constructing consistent multiscale scenarios by transdisciplinary processes: the case of mountain regions facing global change. Ecology and Society 18.

Bryant LD, Hsu-Kim H, Gantzer PA, and Little JC. 2011. Solving the problem at the source: Controlling Mn release at the sediment-water interface via hypolimnetic oxygenation. Water Research 45: 6381–92.

Caraco NF, Cole JJ, and Strayer DL. 2006. Top down control from the bottom: regulation of eutrophication in a large river by benthic grazing. Limnology and Oceanography 51: 664–670.

Cardinale BJ, Duffy JE, Gonzalez A, et al. 2012. Biodiversity loss and its impact on humanity. Nature 486: 59.

Carpenter S, Walker B, Anderies JM, and Abel N. 2001. From metaphor to measurement: resilience of what to what? Ecosystems 4: 765–781.

Chapin III FS, Carpenter SR, Kofinas GP, et al. 2010. Ecosystem stewardship: sustainability strategies for a rapidly changing planet. Trends in ecology & evolution 25: 241–249.

Conversi A, Dakos V, G\aardmark A, et al. 2015. A holistic view of marine regime shifts. Phil Trans R Soc B 370: 20130279.

Crawford JW, Deacon L, Grinev D, et al. 2011. Microbial diversity affects self-organization of the soil–microbe system with consequences for function. Journal of the Royal Society Interface: rsif20110679.

De Laender F, Rohr JR, Ashauer R, et al. 2016. Reintroducing environmental change drivers in biodiversity–ecosystem functioning research. Trends in ecology & evolution 31: 905–915.

Desjardins E, Barker G, Lindo Z, et al. 2015. Promoting resilience. The Quarterly review of biology 90: 147–165.

Díaz S, Demissew S, Carabias J, et al. 2015. The IPBES Conceptual Framework— connecting nature and people. Current Opinion in Environmental Sustainability 14: 1–16.

Donohue I, Hillebrand H, Montoya JM, et al. 2016. Navigating the complexity of ecological stability. Ecology Letters 19: 1172–85.

Folke C, Carpenter S, Walker B, et al. 2010. Resilience thinking: integrating resilience, adaptability and transformability. Ecology and society 15: https://www.jstor.org/stable/26268226.

Gedan KB, Altieri AH, and Bertness MD. 2011. Uncertain future of New England salt marshes. Marine Ecology Progress Series 434: 229–238.

Griffiths BS and Philippot L. 2013. Insights into the resistance and resilience of the soil microbial community. FEMS microbiology reviews 37: 112–129.

Grimm V and Wissel C. 1997. Babel, or the ecological stability discussions: an inventory and analysis of terminology and a guide for avoiding confusion. Oecologia 109: 323–334.

Ingrisch J and Bahn M. 2018. Towards a comparable quantification of resilience. Trends in Ecology & Evolution 33: 251–259.

Isaac, N.J.B. et al 2018. Defining and delivering resilient ecological networks: Nature conservation in England. Journal of Applied Ecology (in press) doi 10.111/1365-2664.13196.

Jax K, Barton DN, Chan KM, et al. 2013. Ecosystem services and ethics. Ecological Economics 93: 260–268.

Jax K and Heink U. 2015. Searching for the place of biodiversity in the ecosystem services discourse. Biological Conservation 191: 198–205.

Kathol M, Fischer H, and Weitere M. 2011. Contribution of biofilm-dwelling consumers to pelagic–benthic coupling in a large river. Freshwater Biology 56: 1160–1172.

Lindner M, Maroschek M, Netherer S, et al. 2010. Climate change impacts, adaptive capacity, and vulnerability of European forest ecosystems. Forest ecology and management 259: 698–709.

Mace GM, Norris K, and Fitter AH. 2012. Biodiversity and ecosystem services: a multilayered relationship. Trends in ecology & evolution 27: 19–26.

Mayer PM, Reynolds SK, McCutchen MD, and Canfield TJ. 2007. Meta-Analysis of Nitrogen Removal in Riparian Buffers. Journal of Environmental Quality 36: 1172–80.

McCay DPF, Peterson CH, DeAlteris JT, and Catena J. 2003. Restoration that targets function as opposed to structure: replacing lost bivalve production and filtration. Marine Ecology Progress Series 264: 197–212.

Mehner T, Diekmann M, Gonsiorczyk T, et al. 2008. Rapid recovery from eutrophication of a stratified lake by disruption of internal nutrient load. Ecosystems 11: 1142–1156.

Millennium Ecosystem Assessment MEA. 2005. Ecosystems and human well-being. Island Press Washington, DC.

Minchin D, Lucy F, and Sullivan M. 2002. Zebra mussel: impacts and spread. In: Invasive aquatic species of Europe. Distribution, impacts and management. Springer.

Newton AC. 2016. Biodiversity risks of adopting resilience as a policy goal. Conservation Letters 9: 369–376.

Oliver TH, Heard MS, Isaac NJB, et al. 2015. Biodiversity and Resilience of Ecosystem Functions. Trends in Ecology & Evolution xx: 1–12.

Palumbi SR, Sandifer PA, Allan JD, et al. 2009. Managing for ocean biodiversity to sustain marine ecosystem services. Frontiers in Ecology and the Environment 7: 204–211.

Parry M, Canziani O, Palutikof J, et al. 2007. Climate change 2007: impacts, adaptation and vulnerability. Cambridge University Press Cambridge.

Peterson GD, Cumming GS, and Carpenter SR. 2003. Scenario planning: a tool for conservation in an uncertain world. Conservation biology 17: 358–366.

Polasky S, Carpenter SR, Folke C, and Keeler B. 2011. Decision-making under great uncertainty: environmental management in an era of global change. Trends in ecology & evolution 26: 398–404.

Ponge J-F, Pérès G, Guernion M, et al. 2013. The impact of agricultural practices on soil biota: a regional study. Soil Biology and Biochemistry 67: 271–284.

Rode M, Hartwig M, Wagenschein D, et al. 2015. The importance of hyporheic zone processes on ecological functioning and solute transport of streams and rivers. In: Ecosystem services and river basin ecohydrology. Springer.

Sasaki T, Furukawa T, Iwasaki Y, et al. 2015. Perspectives for ecosystem management based on ecosystem resilience and ecological thresholds against multiple and stochastic disturbances. Ecological indicators 57: 395–408.

Schimel J and Schaeffer SM. 2012. Microbial control over carbon cycling in soil. Frontiers in microbiology 3: 348.

Schmidt MW, Torn MS, Abiven S, et al. 2011. Persistence of soil organic matter as an ecosystem property. Nature 478: 49.

Schoon ML, Robards MD, Brown K, et al. 2015. Politics and the resilience of ecosystem services. Principles for building resilience: sustaining ecosystem services in social-ecological systems Cambridge University Press, Cambridge, UK http://dx.doi.org/101017/cbo9781316014240 3: 32–49.

Seidl R, Spies TA, Peterson DL, et al. 2016. Searching for resilience: addressing the impacts of changing disturbance regimes on forest ecosystem services. Journal of Applied Ecology 53: 120–129.

Seppelt R, Dormann CF, Eppink FV, et al. 2011. A quantitative review of ecosystem service studies: approaches, shortcomings and the road ahead. Journal of applied Ecology 48: 630–636.

Spears BM, Ives SC, Angeler DG, et al. 2015. Effective management of ecological resilience–are we there yet? Journal of applied ecology 52: 1311–1315.

Standish RJ, Hobbs RJ, Mayfield MM, et al. 2014. Resilience in ecology: Abstraction, distraction, or where the action is? Biological Conservation 177: 43–51.

Strayer DL and Smith LC. 1996. Relationships between zebra mussels (Dreissena polymorpha) and unionid clams during the early stages of the zebra mussel invasion of the Hudson River. Freshwater Biology 36: 771–780.

Thompson ID, Okabe K, Parrotta JA, et al. 2014. Biodiversity and ecosystem services: lessons from nature to improve management of planted forests for REDD-plus. Biodiversity and conservation 23: 2613–2635.

Timpane-Padgham BL, Beechie T, and Klinger T. 2017. A systematic review of ecological attributes that confer resilience to climate change in environmental restoration. PLoS One 12: e0173812, https://doi.org/10.1371/journal.pone.0173812.

Traill LW, Lim ML, Sodhi NS, and Bradshaw CJ. 2010. Mechanisms driving change: altered species interactions and ecosystem function through global warming. Journal of Animal Ecology 79: 937–947.

Wermelinger B. 2004. Ecology and management of the spruce bark beetle Ips typographus— a review of recent research. Forest ecology and management 202: 67–82.

