## Supplement for "Resilience trinity: safeguarding ecosystem services across three different time horizons and decision contexts"

Table S1: Resilience mechanisms described in literature and definitions given by the authors.

| Mechanism | Reference | Definition |
|---|---|---|
| **Adaptive capacity** | Bernhardt and Leslie 2013 | “ability of populations, communities and ecosystems to adapt […] through a combination of phenotypic plasticity, physiol. responses, distributional shifts, rapid evolution of traits” |
| **Adaptive phenotypic plasticity** | Oliver et al. 2015 | “capacity of individuals to respond to environmental changes through flexible behavioral or physiological strategies […]” |
| **Area of natural habitat cover at the landscape scale** | Oliver et al. 2015 | “larger areas of natural or seminatural habitat tend to provide a greater range and amount of ressources, which promote higher spec. richness and larger population sizes […]. This […] is likely to mean greater genetic diversity and functional redundancy […]” |
| **Biodiversity** | Palumbi et al. 2009, Chapin et al. 2010, Griffiths and Philippot 2013, Thompson et al. 2014 | “insurance hypothesis […] based on the intuitive idea that the probability of finding species able to adapt to adapt to changing conditions […] is greater in a more diverse ecosystem”) (Griffiths and Philippot 2013) |

| Mechanism | Reference | Definition |
|---|---|---|
| **Connectivity** | Biggs et al. 2012, Bernhardt and Leslie 2013 | “connections that promote stability and recovery at multiple scales of biological organization” (Bernhardt and Leslie 2013) |
| **Dispersal ability** | Bernhardt and Leslie 2013 | “A species` ability to expand its range into more […] suitable habitats“ |
| **Diversity** | Chapin et al. 2010, Biggs et al. 2012, Bernhardt and Leslie 2013, Desjardins et al. 2015 | “Diversity among elements contributing to a particular ES can modify the effects of disturbance itself” (Biggs et al. 2012) |
| **Dominant species** | Sasaki et al. 2015 | “if the dominant species is resilient to disturbances” it will maintain ES functioning despite disturbances |
| **Negative feedbacks** | Chapin et al. 2010, Gedan et al. 2011, Biggs et al. 2012, Conversi et al. 2014, Spears et al. 2015 | “Negative feedback mechanisms contribute to maintain ecosystem the ecosystem state” (Conversi et al. 2014) |
| **Functional diversity** | Chapin et al. 2010 | “As with economic systems, and ecosystem whose species have a narrow range of functional properties […] has a limited capacity to adjust to change and to sustain ecosystem services (functional diversity) compared with an ecosystem with greater functional and response diversity.” |

| Mechanism | Reference | Definition |
|---|---|---|
| **Functional redundancy** | Biggs et al. 2012, Bernhardt and Leslie 2013, Griffiths and Philippot 2013, Desjardins et al. 2015, Oliver et al. 2015, Alexander et al. 2016 | "when multiple species perform similar functions (…) the resistance of an ecosystem function will be higher if those species also have differing responses to environmental perturbations" (Oliver et al. 2015) |
| **Genetic diversity** | Palumbi et al. 2009, Thompson et al. 2014 | "Resilience is an emergent ecosystem property conferred through biodiversity, related to genetic diversity, species diversity (…) and ecosystem diversity" (Thompson et al. 2014) |
| **Genetic variability** | Oliver et al. 2015 | "Higher adaptive genetic variation increases the likelihood that genotypes that are tolerant to a given environmental perturbation will be present in a population" |
| **Habitat diversity** | Palumbi et al. 2009 | - |
| **Individual response** | Griffiths and Philippot 2013 | "Response of individual cells to disturbance which has consequences for the stability of the total community" |
| **Intrinsic rate of population increase** | Oliver et al. 2015 | "Species with a high intrinsic rate of increase will recover more quickly from environmental perturbations or show resistance of this population reinforcement occurs during the perturbation" |
| **Keystone species** | Traill et al. 2010, Sasaki et al. 2015 | "loss of the keystone species can lead to cascading effects" (Sasaki et al. 2015) |

| Mechanism | Reference | Definition |
|---|---|---|
| **Landscape-heterogeneity** | Thompson et al. 2014, Desjardins et al. 2015 | “Resilience is an emergent ecosystem property conferred through biodiversity, (…), ecosystem diversity (heterogeneity and beta diversity) across a forest landscape“ (Thompson et al 2014) |
| **Landscape-level functional connectivity** | Oliver et al. 2015 | “Metapopulation theory suggests that populations in well-connected landscapes will persist better or recolonize more rapidly (….) (the “rescue” effect)” |
| **Learning** | Chapin et al. 2010, Biggs et al. 2012 | “The process of modifying existing or acquiring new knowledge, behaviors, skills, values, or preferences at individual, group, or societal levels” (Biggs et al. 2012) |
| **Local environmental heterogeneity** | Oliver et al. 2015 | “spatial heterogeneity can enhance the resistence of ecosystem functions by:<br>(i) facilitating the persistence of individual species under environmental perturbations by providing a range of resources and microclimatic refugia; and (ii) increasing overall species richness and, therefore, functional redundancy.” |
| **Modularity** | Bernhardt and Leslie 2013 | “It refers to compartimentalization of populations in space and time. (…).<br>For example, where populations are too closely connected, severe disturbances to one population may affect all populations.” |

| Mechanism | Reference | Definition |
|---|---|---|
| **Network architecture** | Griffiths and Philippot 2013 | "A highly connected and nested architecture promotes community stability in mutualistic networks, whereas stability is increased in compartmented and weakly connected architectures in trophic networks. (…) " |
| **Network interaction structure** | Oliver et al. 2015 | "In general, highly connected nested networks dominated by generalized interactions are less susceptible to cascading extinction effects and provide more resistant ecosystem functions." |
| **Number of links in the food web** | Bernhardt and Leslie 2013 | "Ecosystems with few links are extremely sensitive to the removal of any given species, and many secondary extinctions may result. (…) in highly connected food webs the onset of secondary extinction is delayed." |
| **Participation** | Biggs et al. 2012 | "active engagement of relevant stakeholders in SES management and governance" "Participation appears central to facilitating the collective action required to respond to disturbance (…)" |
| **Polycentric governance systems** | Chapin et al. 2010, Biggs et al. 2012 | "a governance system with multiple, nested governing authorities at different scales." "Policentric structures confer modularity and functional redundancy"(Biggs et al. 2012) |

| Mechanism | Reference | Definition |
|---|---|---|
| **Response diversity** | Chapin et al. 2010, Bernhardt and Leslie 2013, Sasaki et al. 2015 | "the diversity of responses to environmental change within and among species contributing to the same ecosystem function" (Elmqvist et al. 2003 cited in Bernhardt and Leslie 2013) |
| **Sensitivity to environmental change** | Oliver et al. 2015 | "Individuals with [response] traits conferring reduced sensitivity to environmental change will confer higher resistance on ecosystem functions." |
| **Slow variables** | Bennett et al. 2009, Biggs et al. 2012 | "slow variables are usually related to regulating ecosystem services, and that the strength of regulating services can attenuate the impact of shocks on ecosystems." (Bennett et al. 2009) |
| **Species diversity** | Traill et al. 2010, Thompson et al. 2014 | "As a general rule, ecosystem function is most resilient to change from disturbance where species diversity, or key functional species groups are maintained" (Traill et al. 2010) |
| **Species redundancy** | Thompson et al. 2014 | "Species in the same functional group often show different responses to disturbances (Laliberté et al. 2010), and hence the value of redundancy" |
| **Strength of species interactions** | Bernhardt and Leslie 2013 | "Weakly interacting species stabilize community dynamics by dampening strong, potentially destabilizing consumer-resource interactions and facilitative interactions." |

| Mechanism | Reference | Definition |
|---|---|---|
| **Tolerance of environmental stress and capacity to acclimate** | Bernhardt and Leslie 2013 | “ (…) phenotypic plasticity may be the most important component of adaptive potential (…). ” |
| **Top predators** | Thompson et al. 2014 | “Loss of keystone predators can have large effects for a system through cascading effects of expansion of herbivore populations” |

**Table S 2: Literature selection**

| Source | Step | # articles |
|---|---|---|
| Web of Science Search (05-09-2016) | TS=(resilience AND mechanism* AND ecosystem service*) | 21 |
| Literature | Excluded | 10 |
| | Added | 4 |

Based on our previous literature research we added (Chapin et al. 2010; Biggs et al. 2012; Desjardins et al. 2015; Spears et al. 2015) to our search. We excluded publications that used "mechanism" in a different context than resilience or stated that resilience was an important conservation goal for a specific system but not how it should be conserved.

References:

Alexander S, Aronson J, Whaley O, Lamb D. 2016. The relationship between ecological restoration and the ecosystem services concept. Ecology and Society 21.

Bennett EM, Peterson GD, Gordon LJ. 2009. Understanding relationships among multiple ecosystem services. Ecology letters 12:1394-1404.

Bernhardt JR, Leslie HM. 2013. Resilience to climate change in coastal marine ecosystems. Annual Review of Marine Science 5:371-392.

Biggs R, et al. 2012. Toward Principles for Enhancing the Resilience of Ecosystem Services. Annual Review of Environment and Resources 37:421-448.

Chapin FS, 3rd, et al. 2010. Ecosystem stewardship: sustainability strategies for a rapidly changing planet. Trends in ecology & evolution 25:241-249.

Conversi A, et al. 2014. A holistic view of marine regime shifts. Philosophical Transactions of the Royal Society B 370:20130279-20130279.

Desjardins E, Barker G, Lindo Z, Dieleman C, Dussault AC. 2015. Promoting Resilience. The Quarterly Review of Biology 90:147-165.

Elmqvist T, Folke C, Nyström M, Peterson G, Bengtsson J, Walker B, Norberg J. 2003. Response diversity, ecosystem change, and resilience. Frontiers in Ecology and the Environment 1:488-494.

Gedan KB, Altieri AH, Bertness MD. 2011. Uncertain future of New England salt marshes. Marine Ecology Progress Series 434:229-237.

Griffiths BS, Philippot L. 2013. Insights into the resistance and resilience of the soil microbial community. FEMS Microbiology Reviews 37:112-129.

Laliberte E, Wells JA, DeClerck F, Metcalfe DJ, Catterall CP, Queiroz C, Aubin I, Bonser SP, Ding Y, Fraterrigo JM. 2010. Land-use intensification reduces functional redundancy and response diversity in plant communities. Ecology letters 13:76-86.

Oliver TH, et al. 2015. Biodiversity and Resilience of Ecosystem Functions. Trends in Ecology and Evolution 30:673-684.

Palumbi SR, et al. 2009. Managing for ocean biodiversity to sustain marine ecosystem services. Frontiers in Ecology and the Environment 7:204-211.

Sasaki T, Furukawa T, Iwasaki Y, Seto M, Mori AS. 2015. Perspectives for ecosystem management based on ecosystem resilience and ecological thresholds against multiple and stochastic disturbances. Ecological Indicators 57:395-408.

Spears BM, Ives SC, Angeler DG, Allen CR, Birk S, Carvalho L, Cavers S, Daunt F, Morton RD, Pocock MJ. 2015. Effective management of ecological resilience–are we there yet? Journal of Applied Ecology 52:1311-1315.

Thompson ID, Okabe K, Parrotta JA, Brockerhoff E, Jactel H, Forrester DI, Taki H. 2014. Biodiversity and ecosystem services: lessons from nature to improve management of planted forests for REDD-plus. Biodiversity and Conservation 23:2613-2635.

Traill LW, Lim ML, Sodhi NS, Bradshaw CJ. 2010. Mechanisms driving change: altered species interactions and ecosystem function through global warming. Journal of Animal Ecology 79:937-947.
